# AAV-mediated CRISPR/Cas gene editing of retinal cells *in vivo*

**DOI:** 10.1101/039156

**Authors:** Sandy SC Hung, Vicki Chrysostomou, Fan Li, Jeremiah KH Lim, Jiang-Hui Wang, Joseph E Powell, Leilei Tu, Maciej Daniszewski, Camden Lo, Raymond C Wong, Jonathan G Crowston, Alice Pébay, Anna E King, Bang V Bui, Guei-Sheung Liu, Alex W Hewitt

## Abstract

**PURPOSE:** CRISPR/Cas has recently been adapted to enable efficient editing of the mammalian genome, opening novel avenues for therapeutic intervention of inherited diseases. In seeking to disrupt Yellow Fluorescent Protein (YFP) in a Thy1-YFP transgenic mouse, we assessed the feasibility of utilising the adeno-associated virus 2 (AAV2) to deliver CRISPR/Cas for gene modification of retinal cells *in vivo*.

**METHODS:** sgRNA plasmids were designed to target *YFP* and after *in vitro* validation, selected guides were cloned into a dual AAV system. One AAV2 construct was used to deliver SpCas9 and the other delivered sgRNA against *YFP* or *LacZ*(control) in the presence of mCherry. Five weeks after intravitreal injection, retinal function was determined using electroretinography and CRISPR/Cas-mediated gene modifications were quantified in retinal flat mounts.

**RESULTS:** AAV2-mediated *in vivo* delivery of SpCas9 with sgRNA targeting *YFP*, significantly reduced the number of YFP fluorescent cells of the inner retina of our transgenic mouse model. Overall, we found an 84.0% (95% CI: 81.8-86.9) reduction of YFP-positive cells in *YFP*-sgRNA infected retinal cells compared to eyes treated with *LacZ*-sgRNA. Electroretinography profiling found no significant alteration in retinal function following AAV2-mediated delivery of CRISPR/Cas components compared to contralateral untreated eyes.

**CONCLUSIONS:** Thy1-YFP transgenic mice were used as a rapid quantifiable means to assess the efficacy of CRISPR/Cas-based retinal gene modification *in vivo*. We demonstrate that genomic modification of cells in the adult retina can be readily achieved by viral mediated delivery of CRISPR/Cas.

## INTRODUCTION

Many ophthalmic diseases manifest due to well-defined genetic mutations, and inherited retinal diseases now comprise the leading cause of blind registrations in working-aged individuals.^1^ Inherited retinal dystrophies are a very heterogenous group of conditions, all of which are as yet currently untreatable.^2,3^ For example, over 100 disease causing variants across 21 loci have been found to cause Leber Congenital Amaurosis.^4^ To-date, much work has focused on viral-mediated gene-replacement therapy, where gene expression is augmented by ectopic replacement of a normal gene product.^5^ However, emerging data suggests that the efficacy of such viral-mediated gene-replacement therapy may reduce over time.^6,7^ Additionally, genetic heterogeneity and restrictions in viral payloads limit the broad application of such an approach for many inherited retinal diseases.^8–10^

The Clustered Regularly-Interspaced Short Palindromic Repeats (CRISPR) and CRISPR-associated protein (Cas) system used by bacteria to counter viral intrusion, has recently been adapted to allow efficient editing of the mammalian nuclear genome.^11,12^ CRISPR/Cas-based technology is particularly attractive for treating inherited diseases caused by genes with very specific spatial and stoichiometric expression^5^.

There have been a small number of studies, which have demonstrated the potential of *in vivo* gene editing for therapeutic applications.^13^ Although the first reported *in vivo* application of CRISPR/Cas used the endogenous homology directed repair pathway to correct the precise disease-causing mutation,^14^ the majority of subsequent reports have used CRISPR/Cas for gene knockout.^15–17^ In postmitotic cells, such as those found in the adult retina, double stranded DNA breaks are preferentially repaired through non-homologous end joining, and as such, diseases caused by gain-of-function, dominant negative or increased copy number variants could be readily amenable to preemptive intervention whereby CRISPR/Cas technology is used to disrupt the mutant allele.

CRISPR/Cas gene editing has been applied to the retina *in vivo* using electroporation.^18,19^ Wang and colleagues electroporated constructs expressing *Streptococcus pyogenes* Cas9 (SpCas9) to disrupt the function of *Blimp1* and thereby dissect the molecular pathways involved in the regulation of murine retinal rod and bipolar cell development *in vivo*.^18^ More recently, Bakondi *et al*. reported the use of electroporation to transfect photoreceptors with a plasmid containing SpCas9 and a single guide RNA (sgRNA) targeting the *Rho* gene in rats harbouring the dominant *Rho* S334ter mutation.^19^ Strikingly, this intervention was found to prevent retinal degeneration and improve visual function.^19^ Nonetheless, despite these promising applications it is clear that such approaches for CRISPR/Cas delivery *in vivo* are not currently applicable as a human therapy.

The aim of our work was to assess the feasibility and efficiency of CRISPR/Cas-mediated gene editing in the retina using a viral delivery system that could be adapted for use in a clinical setting in the future. To objectively quantify a phenotypic outcome for *in vivo* gene editing, we designed sgRNA constructs to disrupt a yellow fluorescent protein (*YFP*) in a transgenic mouse. As such, we report the use of a robust, quantifiable means to explore the efficacy of a CRISPR/Cas system for retinal modification and demonstrate clear proof-of-concept evidence for gene modification *in vivo*.

## METHODS

### Ethics Approval and Colony Maintenance

Ethics approval for this work was obtained from the Animal Ethics Committee of the University of Tasmania (A14827) and the St Vincent’s Hospital (AEC 014/15), in accordance with the requirements of the National Health and Medical Research Council of Australia (Australian Code of Practice for the Care and Use of Animals for Scientific Purposes). We adhered to the ARVO Statement for the Use of Animals in Ophthalmic and Vision Research.

Thy1-YFP transgenic mice [B6.Cg-Tg(Thy1-YFP)16Jrs/J], which express YFP in all retinal ganglion cells (RGCs), as well as amacrine cells and bipolar cells in the retina, were obtained from the Jackson Laboratory (mouse Stock No: 003709) and bred at the mouse facility of the Menzies Institute for Medical Research (Hobart, Australia).^20^ Mice were housed in standard conditions (20°C, 12–12 hours light-dark cycle) with access to food and water *ad libitum*.

### sgRNA Design and Construct Generation

sgRNA targeting the 5’ region of the *YFP* gene were designed using the CRISPR design tool (http://crispr.mit.edu/)(Supplementary Table 1).^21^ The MM9 mouse assembly was used as the reference genome and three sgRNAs were selected for subsequent profile testing. YFP overexpressing 3T3-L1 (ATCC® CL-173; ATCC, Manassas, VA, USA) and HEK293A (Catalogue number R70507; Life Technologies, Mulgrave, Victoria, Australia) cell lines were used for indel analysis and *in vitro* validation, respectively (Figure 1). Control sgRNA sequence targeting *LacZ* was based on the work by Swiech and colleagues.^22^ sgRNAs were initially cloned into the pSpCas9(BB)purov2.0 vector (a gift from Feng Zhang; Addgene plasmid #62988),^21^ and the best performing sgRNAs were subsequently cloned into pX552-mCherry (a gift from Feng Zhang; Addgene plasmids #60957 and #60958).^22^ pX552-mCherry was generated by replacing the *GFP* with *mCherry* using SalI and BspEI restriction sites.

**Figure 1.**
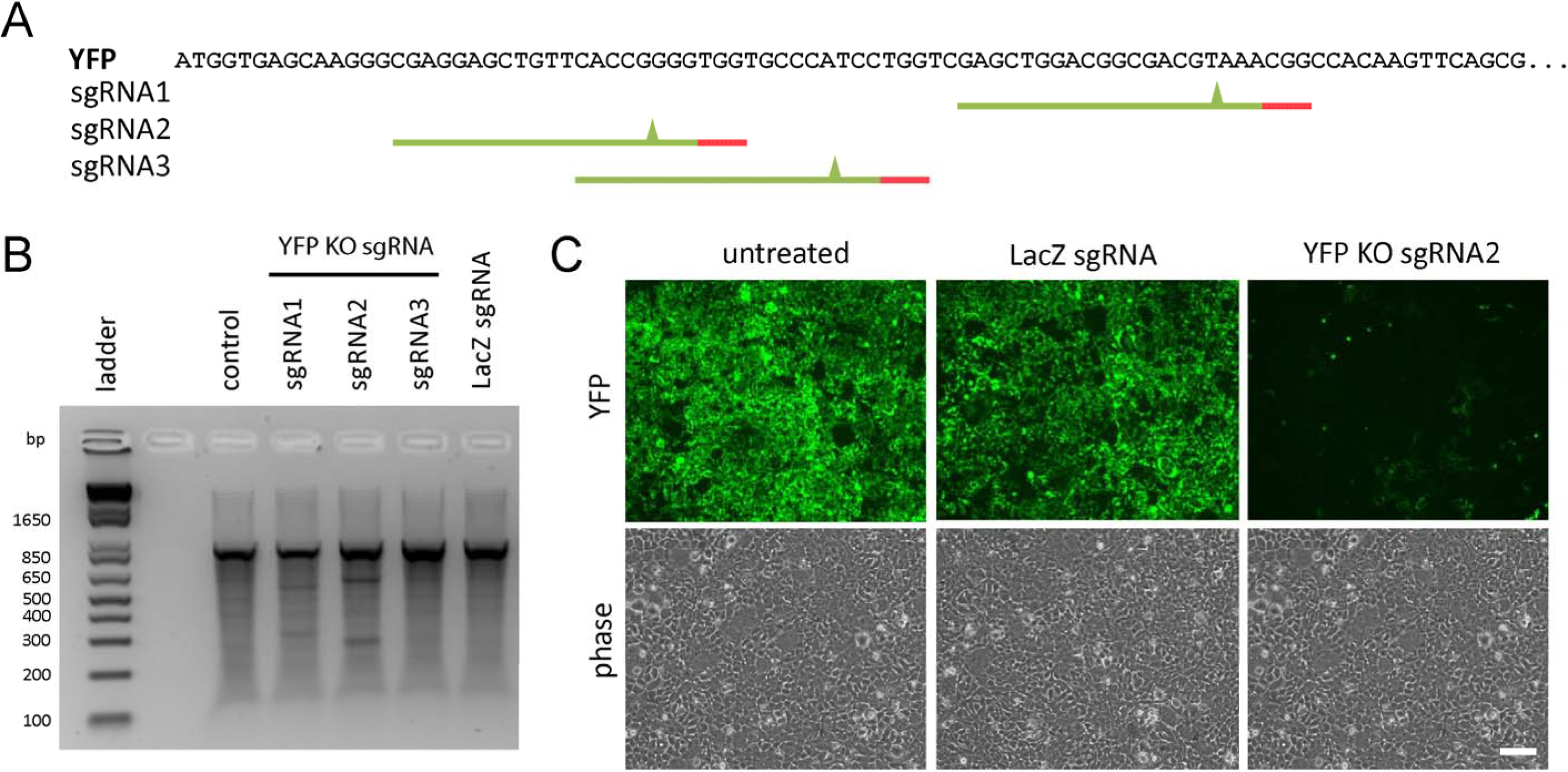
sgRNA design, plasmid construction and validation for knockout (KO) of Yellow-Fluorescent Protein (YFP). (**A**) YFP target sequence used for sgRNA design. Selected sgRNA ideograms display the PAM sequence (red) and putative cut site (green triangle). Indel analysis using a SURVEYOR assay identified “YFP KO sgRNA2” as having the best KO efficiency in our 3T3-L1-YFP line (**B**), and this was confirmed *in vitro* using the HEK293A-YFP line (**C**). Scale bar = 100μm.

### Cell Culture and Transfection

Stable cell lines of HEK293A and mouse 3T3-L1 expressing YFP were generated using pAS2.EYFP.puro lentivirus (RNAiCore, Academia Sinica, Taipei, Taiwan) and selected using puromycin or FACS sorting. All media components were purchased from Life Technologies. Cell lines were maintained in DMEM (Catalogue number 11995040) supplemented with 10% FBS (Catalogue number 26140079), 2mM L-glutamine (Catalogue number 25030081) and 50U/mL Penicillin-Streptomycin (Catalogue number 15070063) and cultured at 37°C with 5% CO_2_ incubation. Transfection of CRISPR/Cas constructs was performed using Lipofectamine 2000 (Catalogue number: 11668027; Life Technologies), with 2.5μg plasmid per well of 6-well plate, according to the manufacturer’s instructions. Fluorescence images were visualised and captured with a Nikon Eclipse TE2000 inverted microscope (Nikon, Melville, NY, USA).

### Indel Analysis

Genomic DNA was extracted using QIAamp DNA mini kit (Catalogue number 51304; Qiagen, Chadstone, Victoria, Australia). Indel analysis was performed with the SURVEYOR assay kit (Catalogue number 706020; Integrated DNA Technologies, Inc. Coralville, USA) using 200-400ng of PCR products generated from genomic DNA of transfected cells. SURVEYOR primers flanking the cleavage sites are listed in Supplementary Table 1. PCR cleaved products were resolved on 1.8% TBE agarose gel and visualised using GelRed nucleic acid gel stain (Catalogue number 41003; Biotium, Hayward, CA, USA). To determine the spectrum and frequency of indels induced by the sgRNA selected for *in vivo* work, 3T3L1-YFP cells were transfected with 1.5μg of each AAV2 plasmids, pX551 and pX552-mCherry with YFP KO sgRNA2, using Fugene HD (Catalogue number E2311; Promega; Madison, WI, USA) following the manufacturer’s protocol. Cells were collected on Day 6 and FACS sorted for mCherry positive cells and genomic DNA extracted using QuickExtract DNA Extraction Solution (Catalogue Number QE09050; Epicentre, Madison, WI, USA) following the heating cycle 62°C for 20min, 68°C for 20min and 98°C for 20min. PCR product was synthesized using Taq DNA polymerase with ThermoPol buffer (Catalogue number M0267S; New England Biolobs, Ipswich, MA, USA). TOPO TA cloning the PCR fragment into pCR2.1-TOPO vector was conducted following manufacturer’s protocol (Catalogue number 45-0641; Life Technologies). Sanger sequencing was used to identify mutations in the *YFP* fragment using the M13 forward primer (−20) included in the kit.

### Virus production

Recombinant AAV2 viruses were produced in HEK293 cells packaging either pX551, containing SpCas9 or pX552-mCherry with the respective sgRNAs, and pseudo-serotyped with the AAV2 capsid (pXX2) as previously outlined.^23^ Viral vectors were purified by AAV2pro® Purification Kit (Catalogue number 6232; Clontech Laboratories, Inc., Mountain View, CA, USA) and vector genomes were titred by real-time quantitative polymerase chain reaction using the following primer sets: (pX551-F: CCGAAGAGGTCGTGAAGAAG, pX551-R: GCCTTATCCAGTTCGCTCAG, pX552-F: TGTGGAAAGGACGAAACACC, pX552-R: TGGTCCTAAAACCCACTTGC) with the SYBR Green Master Mix (Catalogue number 4309155; Life Technologies).

### Intraocular Injection of AAV vectors

A total of 22 adult mice, aged between 14 and 16 weeks, were randomly allocated to receive YFP-sgRNA or LacZ-sgRNA (n=11 per group). Mice were anesthetized by intraperitoneal injection of ketamine (60 mg/kg) and xylazine (10 mg/kg). Intravitreal injections were performed using a surgical microscope similar to that previously described.^24^ In brief, after making a guide track through the conjunctiva and sclera at the superior temporal hemisphere using a 30-gauge needle, a hand-pulled glass micropipette was inserted into the mid-vitreal cavity. A total of 1 μL of dual-viral suspension (AAV2-SpCas9 3x10^9^ vector genomes (vg)/μL with AAV2-LacZ-sgRNA 2.5x10^9^ vg/μL or AAV2-YFP-sgRNA 3x10^9^ vg/μL) or Balanced Salt Solution was injected into each eye at a rate of 100 nL/s using a UMP3-2 Ultra Micro Pump (World Precision Instruments, Inc. Sarasota, USA). Patency was confirmed following needle removal.

### Electrophysiology Assessment and Retinal Morphology

Mice were euthanized five weeks after intraocular injection. Immediately prior to terminal anesthesia, electroretinography was used to assess retinal function in six mice treated with SpCas9/YFP-sgRNA and six mice treated with SpCas9/LacZ-sgRNA.

Our methods for electroretinogram recordings have been well described previously.^25–27^ In brief, overnight dark-adaptation (minimum 12 hours) was performed and all animals were prepared for recording under dim red illumination. Following general anesthesia (intramuscular ketamine 80 mg/kg, xylazine 10 mg/kg), pupil dilation and corneal anesthesia were obtained with 0.5% tropicamide and 0.5% proxymetacaine, respectively. A Ganzfeld sphere (Photometric Solutions International, Oakleigh, Victoria, Australia) was used to deliver luminous energy ranging from –6.26 to 2.07 log c·dm^−2^·s calibrated with an IL1700 integrating photometer (International Light Technologies, Peabody, MA, USA). Electrical signals were recorded with a chlorided silver electrode on the corneal apex and referenced to a ring electrode placed on the conjunctiva posterior to the limbus. A ground needle electrode was placed in the tail. Full-field flash electroretinograms were recorded using standard settings. Response amplitudes of the scotopic a-wave, scotopic b-wave, scotopic threshold response (STR), photopic negative response and photopic b-wave were measured. Signals were amplified (×1000) over a band-pass of 0.3 to 1000 Hz (–3 dB) and digitized using an acquisition rate of 4 kHz. Ganglion cell response was measured as the peak-to-trough amplitude of the STR recorded at –5.25 log cd·m^−2^·s.^25–27^

Retinal morphology was assessed *in vivo* using spectral domain optical coherence tomography (OCT, Bioptigen, Morrisville, NC, USA). Immediately following ERG recordings, a 1.4 mm wide horizontal B-scan (consisting of 1000 A-scans) of the retina centered at the optic nerve head was obtained. Total retinal thickness (inner retinal surface to retinal pigment epithelium), retinal nerve fibre layer (RNFL; inner retinal surface to nerve fibre layer) and ganglion cell complex (GCC) thickness (inner retinal surface to inner plexiform layer) were returned after manually segmenting key retinal layers (in a masked fashion) followed by averaging thicknesses across a region of interest of 200 to 400 μm from centre of the optic nerve. Values from both nasal and temporal retina were averaged.

### Retinal Tissue Processing

Enucleated eyes were fixed in 4% paraformaldehyde (PFA; Catalogue number C004; ProSciTech, Australia) for 1 hour, and retinal flat mounts were prepared and stained with a 4',6-diamidino-2-phenylindole (DAPI; 0.2 ug/mL; Catalogue number D9542; Sigma-Aldrich; Sydney, NSW, Australia) counterstain. For histological assessment, enucleated eyes were fixed in 4% PFA for 1 hour and embedded in optimal cutting temperature compound prior to frozen sectioning on a microtome-cryostat. Serial 10 μm thick sections were cut, mounted on silanated glass slides, and then stained with DAPI.

### Retinal Imaging, Cell Counting and Statistical Analysis

A fluorescence microscope (Zeiss Axio Imager Microscope, Carl-Zeiss-Strasse, Oberkochen, Germany) equipped with a charge-coupled device digital camera (Axiocam MRm, Zeiss) and image acquisition software (ZEN2, Zeiss) was used. Following whole mounting, each retinal quadrant was photographed using an appropriate filter for the fluorescence emission spectra of mCherry (605nm, Zeiss Filter set 64HE) and YFP (495nm, Zeiss Filter set 38HE). Confocal images were taken using a Nikon C1 confocal laser scanning microscope using a 20x 0.75NA lens and images were tile stitched using the NIS Elements AR program (Nikon, Melville, NY, USA). Retinal cell quantification was performed using individual fluorescent images captured at 400x magnification. A total of 18 images from four flat-mounted eyes treated with SpCas9/LacZ-sgRNA, and 26 images from five flat-mounted eyes treated with SpCas9/YFP-sgRNA were quantified manually using using ImageJ v1.49, by an experienced grader (Fan Li), masked to treatment status.^28^ To investigate retinal cellular subpopulations targeted by our AAV2 vectors, five retinal cross-sections from five SpCas9/LacZ-sgRNA treated eyes and five retinal cross-sections from SpCas9/YFP-sgRNA treated eyes were also quantified (at 200x magnification). Sample group decoding was performed only once data were quantified and analyzed.

The efficiency of YFP knockout was determined by the proportion of YFP negative cells among mCherry expressing cells. Specifically, the mean proportion of YFP-knockout for all eyes treated with SpCas9/YFP-sgRNA was calculated by ΔYFP^−ve^/(ΔYFP^−ve^ + YFP^+ve^), where ΔYFP^−ve^ was the number of mCherry transfected cells in SpCas9/YFP-sgRNA treated eyes, which did not express YFP, minus the average number of mCherry expressing cells in SpCas9/LacZ-sgRNA treated eyes, which did not express YFP (YFP^−ve^); and YFP^+ve^ was the number of cells which expressed both YFP and mCherry.

To profile the putative *in vivo* activity of the promoters used to drive SpCas9 and mCherry expression in our plasmids, we interrogated the single cell mouse retinal expression data generated by Macosko *et al*. (GSE63472).^29^ In brief, transcriptomic data from 44,808 retinal cells collected from P14 C57BL/6 mice were analysed. Hierarchical clustering was used to separate cells into distinct clusters (RGCs, amacrine cells, rods etc) and the percentage of cells in each cluster that expressed Mecp1 and Syn1 (i.e. a transcripts per million (TPM) value ≥1) were quantified. The mean value from clusters of the same cell-type were merged.

Comparisons between continuous traits were made using the paired *t*-test, whilst categorical data were analysed using the Chi-square or Fisher exact test. Unless otherwise specified, data are presented as mean ± SD. A total of 14 images were processed twice to determine intra-grader variability. The intraclass correlation coefficient for YFP^+ve^ cell counts was 0.945 (95% CI: 0.842-0.982) and for YFP^−ve^ cell counts was 0.954 (95% CI: 0.867-0.985). Analyses were performed in the R statistical environment (version 3.2.2).

## RESULTS

### sgRNA Selection and In Vitro Validation

We assessed three sgRNAs targeting the *YFP* sequence, together with a LacZ-sgRNA control,^22^ for cleavage efficiency in 3T3-L1-YFP cells (Figure 1A). Indel analysis, for the quantification of sgRNA-induced nucleotide mismatches revealed that one of our sgRNAs (designated “YFP KO sgRNA2”) targeted *YFP* most effectively (Figure 1B). Following TOPO TA cloning we found approximately 73% cleavage activity of this YFP-sgRNA, which was then used for all subsequent experiments ( Supplementary Figure 1). When transfected into HEK293A-YFP cells, our selected YFP-sgRNA resulted in almost complete reduction in yellow fluorescence compared with untransfected and LacZ-sgRNA controls (Figure 1C).

### In Vivo AAV Delivery and Gene Targeting of CRISPR/Cas9 in the Mouse Retina

To evaluate the YFP disruption *in vivo* by CRISPR/Cas-mediated gene-editing, Thy1-YFP mice received a single intravitreal injection of our dual viral suspension of AAV2-SpCas9 and AAV2-sgRNA-mCherry (Figure 2A). In the murine retina, the promoters used for each plasmid (Mecp2 and Syn1 for SpCas9 and mCherry, respectively) have the greatest expression in RGCs and amacrine cells (Supplementary Figure 2). Five weeks following treatment, inspection of retinal whole mount images, captured under a stereomicroscope, revealed a lower number of YFP positive cells from SpCas9/YFP-sgRNA treated eyes compared to SpCas9/LacZ-sgRNA treated (Figure 2) or contralateral control eyes. To quantify the efficiency of YFP disruption following AAV2 delivery of SpCas9 and sgRNA, high magnification flat mount images were obtained and the proportion of YFP positive and negative, mCherry expressing cells was calculated. A marked decrease in the number of YFP positive cells in the inner retina was found in SpCas9/YFP-sgRNA treated eyes, compared SpCas9/LacZ-sgRNA treated eyes. The proportion of YFP/mCherry expressing cells in eyes treated with SpCas9/YFP-sgRNA was 0.106 ± 0.022, compared to SpCas9/LacZ-sgRNA treated eyes where the proportion of YFP/mCherry expressing cells was 0.664 ± 0.050 (Figure 2D). Overall there was an 84.0% (95% CI: 81.8-86.9) reduction of YFP-positive cells in SpCas9/YFP-sgRNA transfected retinal cells compared to eyes treated with SpCas9/LacZ-sgRNA. As expected, there was no significant variation in the number of YFP cells in untreated control eyes of both groups. Similarly, retinal cross-sections showed a clear disruption of YFP expression in the ganglion cell layer (GCL) of the SpCas9/YFP-sgRNA-treated eyes, but not SpCas9/LacZ-sgRNA-treated eyes (Figure 2F&H). AAV2 was found to predominate in the GCL, with approximately a 50% reduction of YFP expressing RGCs (Supplementary Figure 2).

**Figure 2.**
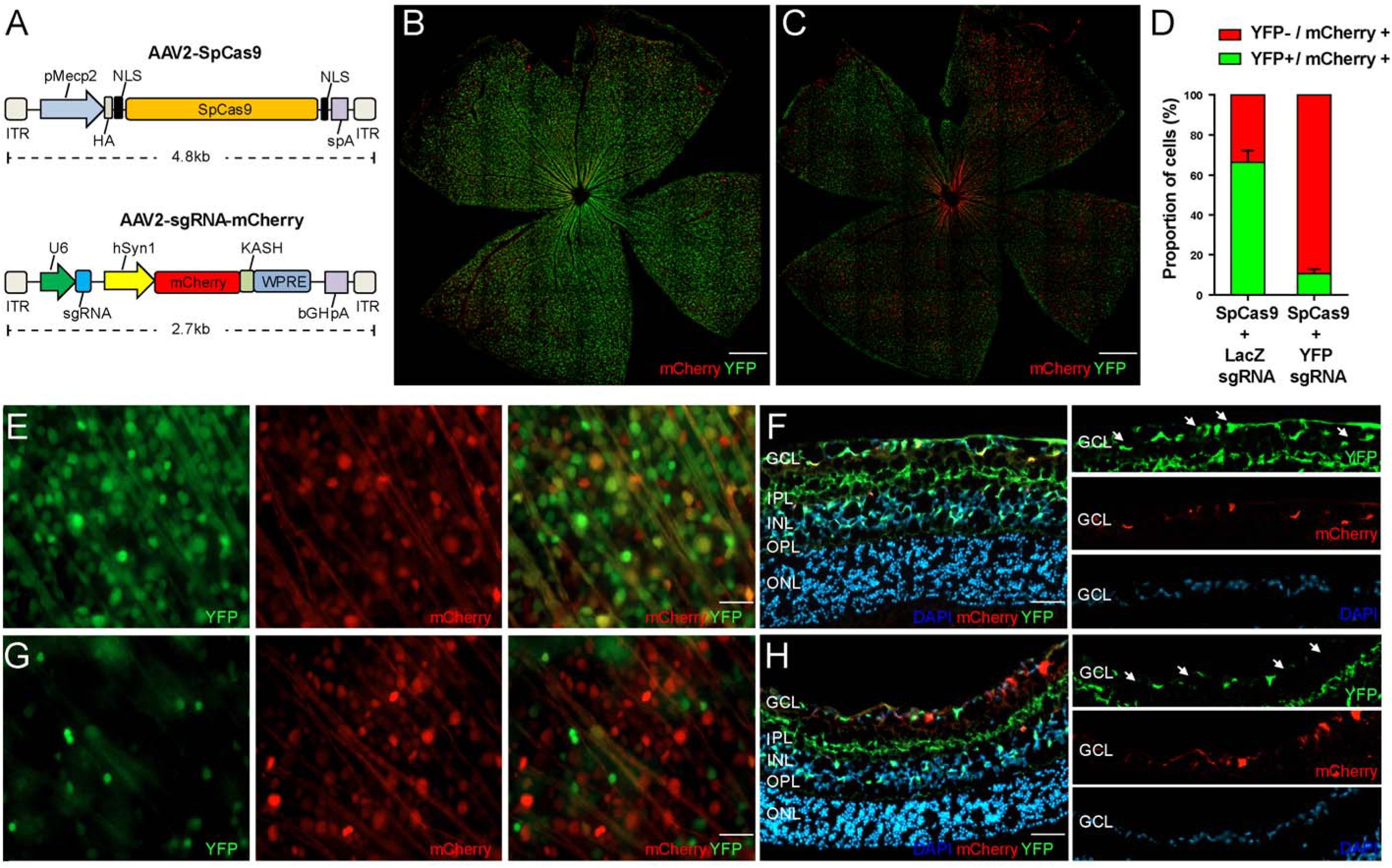
CRISPR/Cas mediated gene-editing of retinal cells *in vivo*. Dual viral suspension of AAV2-SpCas9 and AAV2-sgRNA were used (**A**). sgRNA plasmids also expressed mCherry and the size of the cassettes packaged by AAV2 are displayed. Representative retinal montages from Thy1-Yellow-Fluorescent Protein (YFP) mice exposed *in vivo* to our dual AAV2 plasmid-system carrying SpCas9 and either control (LacZ) sgRNA (**B**) or sgRNAs targeting YFP (**C**) (Scale bar = 500μm). Overall, the proportion of mCherry expressing cells (mCherry+), which lacked YFP (YFP-) was higher in SpCas9/YFP-sgRNA treated eyes (**D**). Higher magnification of flat mount images, displaying differences in YFP expression following AAV2-mediated delivery of SpCas9/LacZ-sgRNA (**E**) or SpCas9/YFP-sgRNA (**G**) (Scale bar = lOμm). Cross sections displaying AAV2 infection of the inner retina confirm YFP knockout in SpCas9/YFP-sgRNA treated eyes (**H**), compared to SpCas9/LacZ-sgRNA treated eyes (**F**) (Scale bar = 50μm). White arrows indicate AAV2-sgRNA infected cells. Abbreviations: GCL, ganglion cell layer; IPL, inner plexiform layer; INL, inner nuclear layer; OPL, outer plexiform layer; ONL, outer nuclear layer; ITR; inverted terminal repeat; pMecp2, truncated methyl-CpG-binding protein 2 promoter; HA, hemagglutinin tag; NLS, nuclear localization signal; spA, synthetic polyadenylation signal; U6, Pol III promoter; sgRNA, single guide RNA; hSyn1, human synapsin 1 promoter; mCherry, monomeric cherry fluorescent protein; KASH, Klarsicht ANC1 Syne Homology nuclear transmembrane domain; WPRE, Woodchuck Hepatitis virus posttranscriptional regulatory element; bGHpA, bovine growth hormone polyadenylation signal.

### Electroretinography Assessment of Retinal Function

Electrophysiological assessment of mice five weeks after treatment with SpCas9/YFP-sgRNA or SpCas9/LacZ-sgRNA, confirmed no adverse effects on photoreceptoral (a-wave: YFP p=0.64; LacZ p=0.35) or bipolar cell (b-wave: YFP p=0.38; LacZ p=0.40) function (Figure 3; Supplementary Figures 3-5). Importantly, given the AAV2 transfection profile following intravitreal injection, inner retinal function as indicated by the oscillatory potentials (YFP p=0.42; lacZ p=0.68) was also not disrupted (Figure 3; Supplementary Figure 3). The scotopic threshold response amplitude, indicative of ganglion cell activity, did not significantly differ between YFP-sgRNA treated and contralateral control eyes (p=0.48), or between LacZ-sgRNA treated and their contralateral eyes (p=0.87)(Supplementary Figure 5).

**Figure 3.**
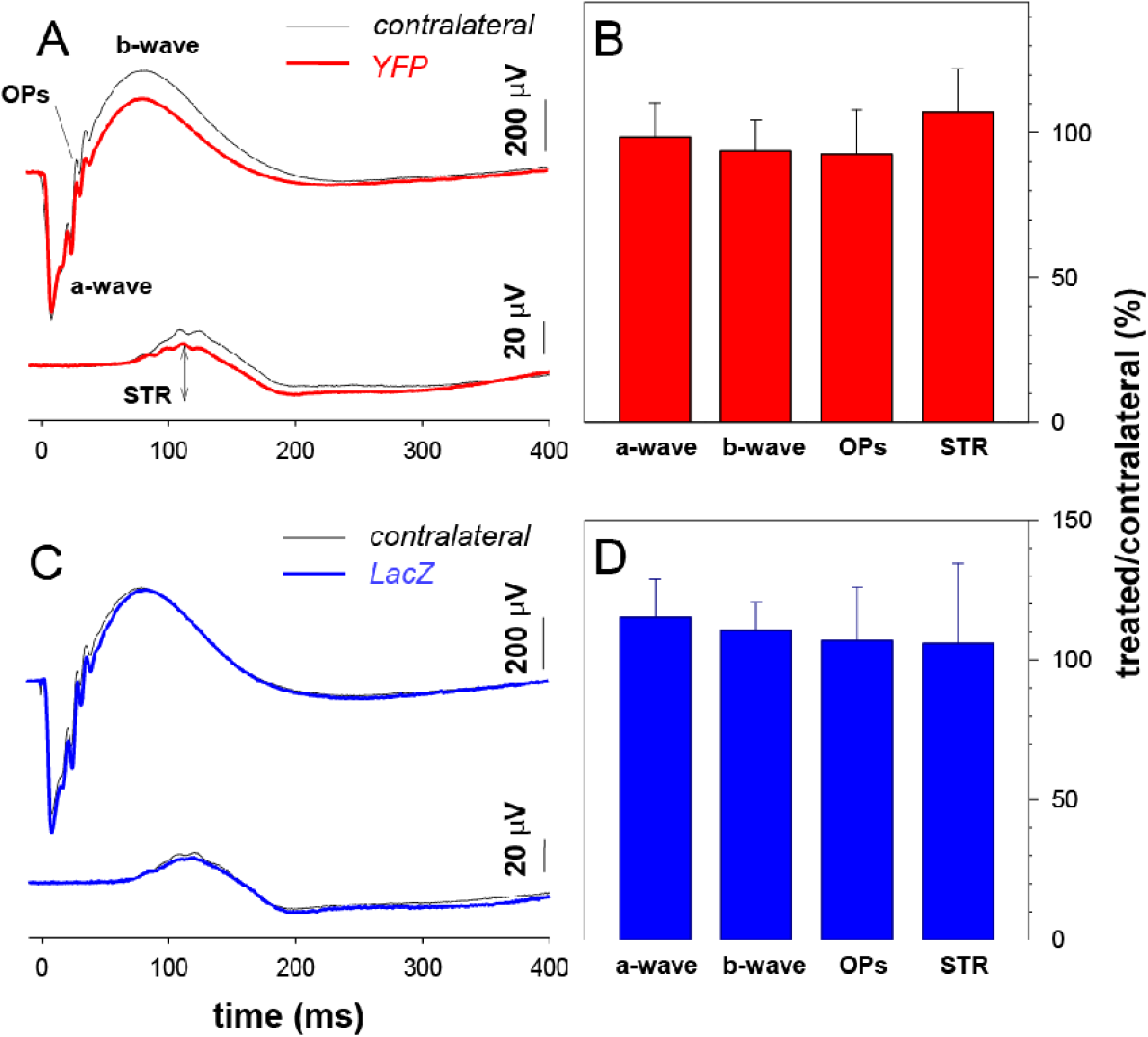
Effect of SpCas9 on retinal function. Averaged ERG waveforms at selected intensities for YFP-sgRNA treated (n=6, red) and contralateral (n=6, black) eyes (**A**). The average photoreceptoral (a-wave), bipolar cell (b-wave), amacrine cell (oscillatory potentials, OPs) and ganglion cell (scotopic threshold response, STR) amplitude in YFP-sgRNA treated relative to contralateral control eyes (%,±SEM) is displayed (**B**). Panel (**C**) displays the averaged ERG waveforms at selected intensities in LacZ-sgRNA treated (n=6, blue) and contralateral eyes (n=6, black). The LacZ-sgRNA group’s average a-wave, b-wave, OPs and STR amplitude relative to contralateral control eyes (%,±SEM)(**D**).

There was no statistically significant difference in retinal morphological profiles *in situ*, between treated or contralateral controls eyes (Supplementary Figure 6). In particular the RNFL and GCC thickness as measured by OCT did not significantly differ between eyes treated with YFP-sgRNA and contralateral control eyes (RNFL p=0.30, GCC p=0.80) or between LacZ-sgRNA and their control eyes (RNFL p=0.70, GCC p=0.39).

## DISCUSSION

We have successfully demonstrated viral-mediated delivery of essential CRISPR/Cas components to retinal cells *in vivo*. Using a transgenic fluorescent mouse model, we have definitive evidence for CRISPR/Cas-mediated gene-editing of *Thy1* expressing retinal cells. With our dual-viral SpCas9/sgRNA suspension we observed a reduction of YFP-positive cells of approximately 84.0% (95% CI: 81.8-86.9). This level of gene modulation *in vivo* is similar to that reported for other tissues, such as brain (~68% using AAV) and liver (80-90% using adenovirus).^22,30^

Our study builds on recent work, where CRISPR/Cas system was introduced into rodent retinas using an electroporation delivery method.^18,19^ By demonstrating that CRISPR/Cas can also cause substantial gene modification activity when introduced by a viral delivery method in the retina, we are closer to translating gene-editing technology for therapeutic purposes. Importantly we found that AAV2-delivered SpCas9 was not retinotoxic over a five week treatment period. Nonetheless, it is clearly possible that adverse effects from ongoing or prolonged CRISPR/Cas expression could arise. As such it will be important to continue to develop CRISPR/Cas systems that could be tightly regulated and thereby be able to create an optimal window for gene modification while reducing the chance for potential off-target activity.^31–34^

AAV vectors have been widely exploited as a promising tool for gene delivery *in vivo* and it is clear that combining alternate AAVs and sgRNA promoters will extend the CRISPR/Cas therapeutic repertoire. Specifically, different AAV serotypes have the ability to transfect distinct cell populations within the retina,^35^ and whilst the AAV2 capsid used in our study is known to primarily transduce retinal ganglion cells when injected intravitreally,^36^ a different AAV2 variant, 7m8, can infect photoreceptors following intravitreal delivery.^37^ Additionally, to overcome the packaging size limitation of AAV vectors,^38^ we used a dual-vector system to package SpCas9 and sgRNA expression cassettes in two separate viral vectors. Although, a single vector that packages a smaller Cas9 ortholog, such as *Staphylococcus aureus* Cas9 (SaCas9), together with a sgRNA, could achieve greater knockout efficiencies *in vivo*,^17,39,40^ dual-vector systems may still be required for mutation correction by enabling the cellular delivery of additional donor templates and appropriate promoter elements. For example, a recent study from Yang and colleagues demonstrated that a two-vector approach with SaCas9 resulted in mutation correction in approximately 10% of hepatocytes in ornithine transcarbamylase-deficient newborn mice.^41^ With ongoing advances in viral engineering and the development of new serotypes, future research investigating the efficacy of different viral-CRISPR/Cas systems in the retina, is warranted.

An important limitation of our work, is the fact that off-target effects were not directly quantified. Nonetheless, to some extent this is a minor consideration for our initial proof-of-concept work which sought to primarily assess “on-target” efficacy of virally delivered CRISPR/Cas in the adult retina. Additionally, it is well appreciated that CRISPR/Cas technology is advancing rapidly and although a major limitation of gene editing has been the prospect of off-target effects, this issue is being addressed. ^33,34^ For example, with careful design of the guide RNA (using a double-nick; truncated sgRNA) it is already possible to avoid most off-target cuts and recently, two independent studies report modifications to SpCas9, which can dramatically reduced off-target activity.^33,34^ Regardless, it is clear that safety will be paramount prior to CRISPR/Cas being used in mainstream medical care, and offtarget profiling will need to be considered before current research is translated into clinical trials.

In summary, this work suggests that a relatively high-efficiency of gene editing in the retina can be achieved by a dual AAV2-mediated CRISPR/Cas system. Whilst *ex vivo* TALEN-based approaches are now in clinical trials, given the versatility and relative ease of design, CRISPR/Cas techniques appear to be more clinically applicable. Inherited eye diseases share several features that make them appealing for the application of *in vivo* gene editing. For example, non-syndromic retinal diseases generally have a specific and well-defined pathogenesis, whereby only a specific subpopulation of cells would need to be targeted. Additionally, following any intervention, the eye can be directly examined for the development of any adverse effects. Nonetheless, any *in vivo* gene editing therapies will require a number of functionally active cells, and it is possible that the threshold for viable cells will vary across genes and diseases.

## ACKNOWLEDGEMENTS

We are grateful to Kenneth Pang, Sze Woei Ng, Elsa Chan, Stacey Jackson, Zheng He and Olivia Swann. This work was supported by grants from the BrightFocus Foundation, the Ophthalmic Research Institute of Australia, Retina Australia, the Childhood Eye Cancer Trust and the Eye Research Australia Fund. AP and BVB are supported by Australian Research Council Future Fellowships. AWH is supported by a National Health and Medical Research Council Practitioner Fellowship. CERA receives operational infrastructure support from the Victorian Government.

